# Gut Microbial succession during conventionalization of germfree chicken

**DOI:** 10.1101/360784

**Authors:** Milton Thomas, Supapit Wongkuna, Sudeep Ghimire, Kinchel C. Doerner, Aaron Singery, Eric Nelson, Tofuko Woyengo, Surang Chankhamhaengdecha, Tavan Janvilisri, Joy Scaria

## Abstract

A gnotobiotic chicken model was developed to study the succession of intestinal microflora from hatching to 18 days of age. Intestinal samples were collected from a local population of feral chickens and administered orally to germ-free 3 day old chicks. Animals were enthanized on 0, 9 and 18 days of age and intestinal samples were collected and subjected to genomic analysis. The five most prevalent phyla were Bacteroidetes (45.73±3.35%), Firmicutes (36.47±2.60%), Proteobacteria (8.28±0.91%), Actinobacteria (5.09±0.52%), and Spriochetes (2.10±0.38%). Principle coordinate analysis indicated the 0, 9 day and 18 day variables clustered together and the microbial communities changed temporally. The Morista-Horn index values ranged from 0.72 to 1, indicating the communities at 0, 9 or 18 days were more similar than dissimilar. The predicted functional profiles of the microbiomes of 0, 9 and 18 days were also similar. These results indicate the gnotobiotic chicks stably maintain the phylogentic diversity and predicted metabolic functionality of the inoculum community.

**Importance:** The domestic chicken is the cornerstone of animal agriculture worldwide with a flock population exceeding 40 billion birds/year. It serves as the economically valuable source of protein globally. Microbiome of poultry has important effects on chicken growth, feed conversion, immune status and pathogen resistance. The significance of our research is in developing a gnotobiotic chicken model to study chicken gut microbiota function. Our experimental model shows that young germfree chicks are able to colonize diverse set of gut bacteria. Therefore, besides using this model to study mechanisms of gut microbiota interactions in the chicken gut, our model could be also used for applied aspects such as determining the safety and efficacy of new probiotic strains derived from chicken gut microbiota.

## Introduction

The chicken gut microbiota influences nutrient utilization (1, 2), immune development (3), endocrine activity (4), development of gastrointestinal tract (5), and detoxification, thus contributing to the improved performance of the birds. The chicken gastrointestinal tract harbors complex communities of bacteria (6, 7). However, cecum harbors the most number of commensals and contains up to 10^11^/ g organisms (8) and therefore was widely studied. In addition to commensal bacteria, cecum also could harbor enteric pathogens that pose both avian and zoonotic health risk (9). The commensals could prevent the colonization of pathogens by competitive exclusion (10) and through the production of bacteriocins (11, 12).

Several experiments were conducted previously to study the microbial succession in broiler chicken intestinal tract (7, 13-15). Also, studies were performed to determine the effect of gut microbes on feed utilization and conversion (1, 16). However, these experiments used 16s RNA sequencing or culture-based techniques. The 16S rRNA sequencing is inherently limited due to bias introduced during PCR reactions. Also, the data has lower resolution and is less efficient in predicting the functional properties of the microbiome. The accuracy of culture-based enumeration of the bacterial population is negatively affected by the inability to grow all the bacteria in culture conditions. In these respect, shotgun metagenomics provides a comprehensive representation of both taxonomical and functional properties of the microbiome. The two studies that used shotgun metagenomics for analyzing chicken microbiome were limited by the number of birds used in those experiments (6, 17).

Feral chickens are derived from domestic chickens that are released to wild and survive many generations. Living in the wilderness induces differences in the feeding habits and social behavioral patterns. Previous research in wild fowls and turkeys have shown that the microbial communities in these birds vary considerably from the domesticated counterparts (18–21). Similar to these findings, we hypothesized that adult feral microbiome could be substantially diverse from the microbial population of the commercial poultry. Use of feral chicken microbiome as probiotic in commercial poultry practices could increase the microbial diversity and thereby possibly provide colonization resistance against enteric pathogens.

The objective of this experiment was to analyze the microbial succession in gnotobiotic chickens when inoculated with adult feral chicken microbiome using shotgun metagenomics. Our findings suggested that feral chicken microbiome could colonize successfully in the young chicken gut without causing detrimental health effects to the host.

## Results

### Presence of *Salmonella* in the Feral Chicken Gut and Sterility of the Isolator

The donor material was collected anaerobically from the cecum and colon contents of 6 feral chickens and immediately was screened for the presence of *Salmonella* by streaking on XLT4 agar plates. All plates were negative for *Salmonella* growth after 24 h incubation at 37°C indicating *Salmonella* was not present in the gut of the feral chickens. The sterility of the isolator and chicks were examined by culturing fecal droppings and swabs from gnotobiotic isolators and found to be negative for bacterial colonies indicating the isolator and chicks were germ-free.

### Phylogenetic distribution of microbiome in gnotobiotic chicken gut compared to inoculum

#### Microbiome composition in the inoculum and cecal contents at phylum level

A taxonomical abundance table with phyla-level distribution was generated in MG-RAST using RefSeq database. The five major phyla in all the samples were Bacteroidetes, Firmicutes, Proteobacteria, Actinobacteria, and Spirochetes (Figure 1). The proportion of Bacteroidetes in the feral chicken inoculum was 66.42% but was lower in 9 d (47.49 ± 4.38%) and 18 d samples (45.73 ± 3.35%). There was no significant difference between the 9 d and 18 d samples for Bacteroidetes. The abundance of Firmicutes in the inoculum was 18% and increased to 27.24 ± 3.88 % in 9 d and to 36.47 ± 2.60% in 18 d samples. The Firmicutes abundance in the chicken gut significantly increased by 18 d compared to 9 d (P > 0.05). The 9 d samples showed a major shift in the proportion of Proteobacteria which nearly doubled compared to inoculum (9.65 % in inoculum vs 16.64 ± 2.78 in 9 d chicken). However, the Proteobacteria level decreased to that in the inoculum by 18 d (8.28 ± 0.91%) and was statistically lower than 9 d samples (P > 0.05). This is in concurrence with the previous findings that facultative anaerobes proliferate initially in the infant gut followed by the outgrowth of the obligate anaerobic bacterial community due to oxygen depletion in the gut (22). Similar to Firmicutes, the abundance of Actinobacteria also increased temporally. In the inoculum, the fraction of Actinobacteria was 1.62%, whereas, in 9 d and 18 d samples, the abundance increased to 3.99 ± 1.41 and 5.09 ± 0.52 % respectively. The percentages of Spirochetes remained similar in the inoculum and gnotobiotic chicken samples at 9 d and 18 d (1.52, 2.58 ± 0.94, and 2.10 ± 0.38%).

**Figure 1.**
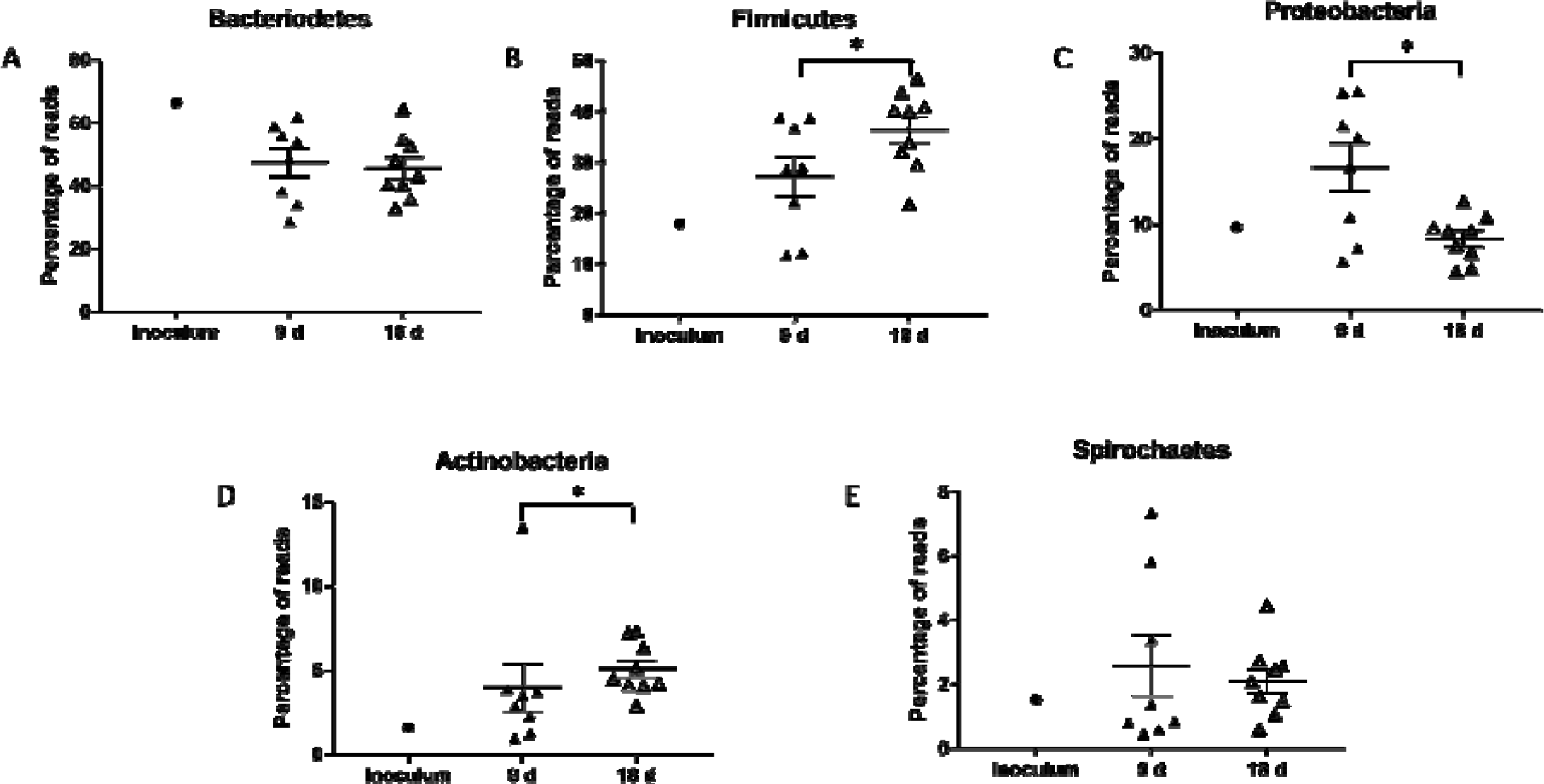
Taxonomical distribution of the major phyla in the inoculum and gnotobiotic chicken gut at 9d and 18 d. Inoculum was derived from 6 healthy feral chicken. Germ-free chicks were inoculated on 3 d post-hatch and euthanized on 9 d (n = 8) and 18 d (n = 9) post-hatch. Cecal contents were collected for DNA isolation. DNA was sequenced using the Miseq 2−250 paired-end sequencing platform. Phylogenetic tables were generated in MG-RAST. The statistical difference in phyla distribution between 9 d and 18 d samples was calculated using non-parametric Mann-Whitney U test. A) Bacteroidetes B) Firmicutes C) Proteobacteria D) Actinobacteria and E) Spirochetes. * represents P > 0.05.

#### Microbiome composition in the inoculum and cecal contents at genera level

At the genus level, the inoculum, 9 d, and 18 d samples was composed predominantly of *Bacteroides* (Figure 2). However, the abundance was higher in the inoculum (52.85 %) compared to 9 d (mean ± SEM; 37.87 ± 3.09 %) and 18 d (35.52 ± 2.53 %) samples. *Clostridium* increased in the 9 d (7.22 ± 0.99 %) and 18 d (9.72 ± 0.74 %) cecal contents compared to inoculum (5.16 %). The next abundant genus was *Prevotella* with the inoculum and gnotobiotic chicken samples indicating similar percentages. *Escherichia*, a member of the Proteobacteria, had higher abundance in the 9 d samples (7.39 ± 1.69%) compared to inoculum (2.28%) and then decreased by 18 d (1.03 ± 0.14%). *Parabacteroides* which represented 4.23% in the inoculum also decreased, similar to *Bacteroides*, to 2.26 ± 0.25 and 2.40 ± 0.18% in 9 d and 18 d samples respectively. The other two major genera that increased in the cecal contents of gnotobiotic chicken compared to inoculum were *Ruminococcus* and *Eubacterium* which are members of Firmicute phylum.

**Figure 2.**
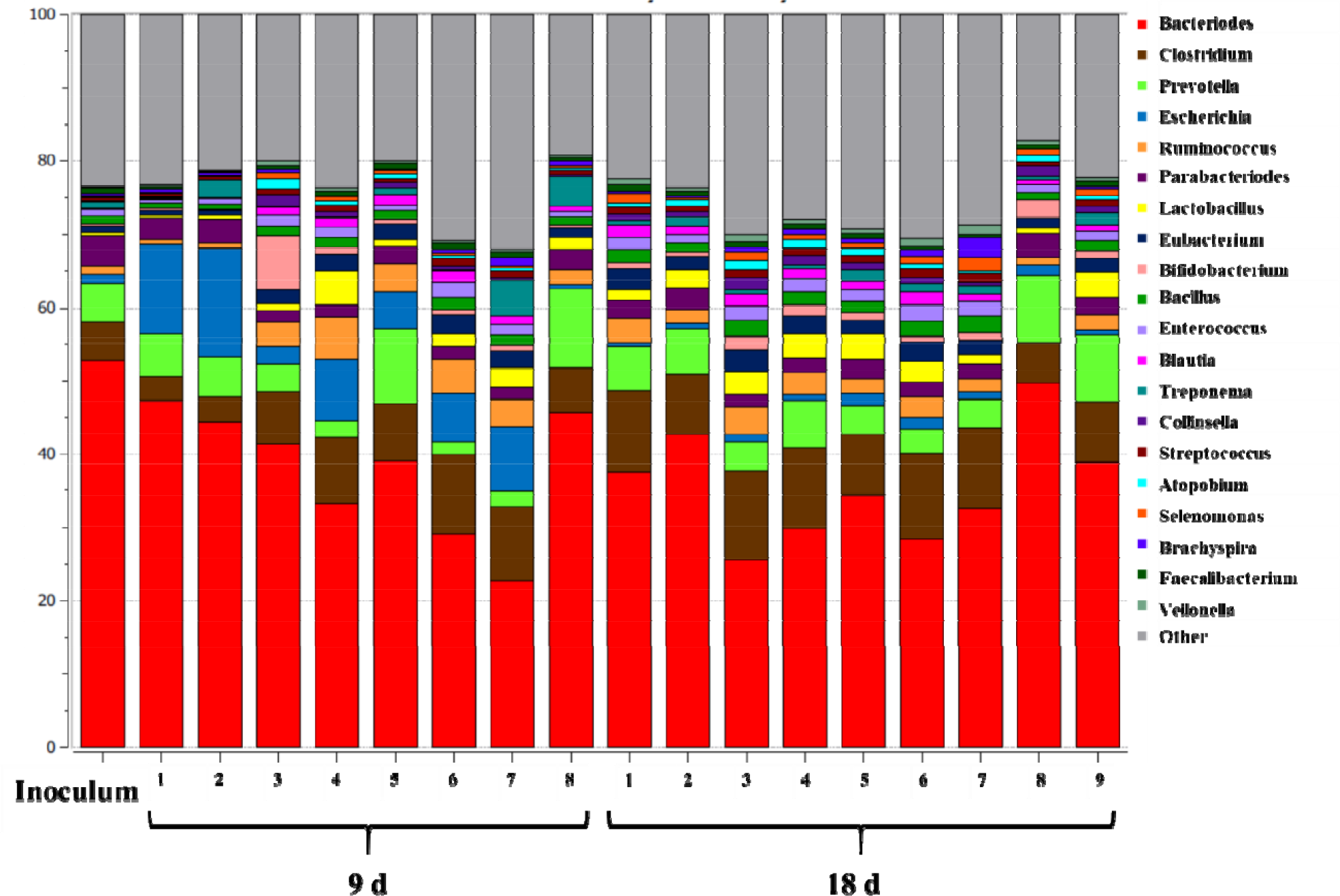
Genera-level distribution of gut microbiome in the gnotobiotic chicken inoculated with intestinal material from feral chickens. The pooled inoculum, derived from 6 healthy feral chicken was orally inoculated to gnotobiotic chicken on 3 d after hatch. Birds were euthanized on 9 d (n = 8) and 18 d (n = 9) of age and cecal contents were collected for DNA isolation. The metagenomic functional analysis was performed in MG-RAST using Refseq database with maximum e-value at 10^−5^ value, minimum percentage identity at 60 %, minimum alignment length of 30 amino acids and abundance 50. Phylogenetic tables were generated in MG-RAST and analysis was conducted using Explicet software.

### Principle coordinate analysis (*PCA*) and β diversity

The principal coordinate analysis (PCA) was calculated using euclidean distance as the similarity metric for clustering the metagenomes (Figure 3 A). While the 9 d communities were randomly distributed across space, the 18 d communities were clustered together and separate from the control, which represented 70.6% of the variation and indicated the microbial communities evolved and matured temporally and attained a similar community profile. These findings are similar to the microbial succession occurring in a previously uninhabited environment such as infant gut where the microbial community attains maturity and stability in the initial years of life (23). Also, the community assemblage in the chicken gut by 18 d resembled closely with the inoculum when compared to 9 d. Genera that were significantly altered in proportion between 9 d and 18 d samples are given in Figure 3 B. *Shigella* and *Escherichia* each decreased by 18 d compared to 9 d. Alternatively, the proportion of *Selenomonas* spp., *Geobacillus* spp., and *Mitsuokella* spp. increased by 18 d. The read percentages of other bacteria that significantly increased by 18 d represented less than 0.1 %.

**Figure 3.**
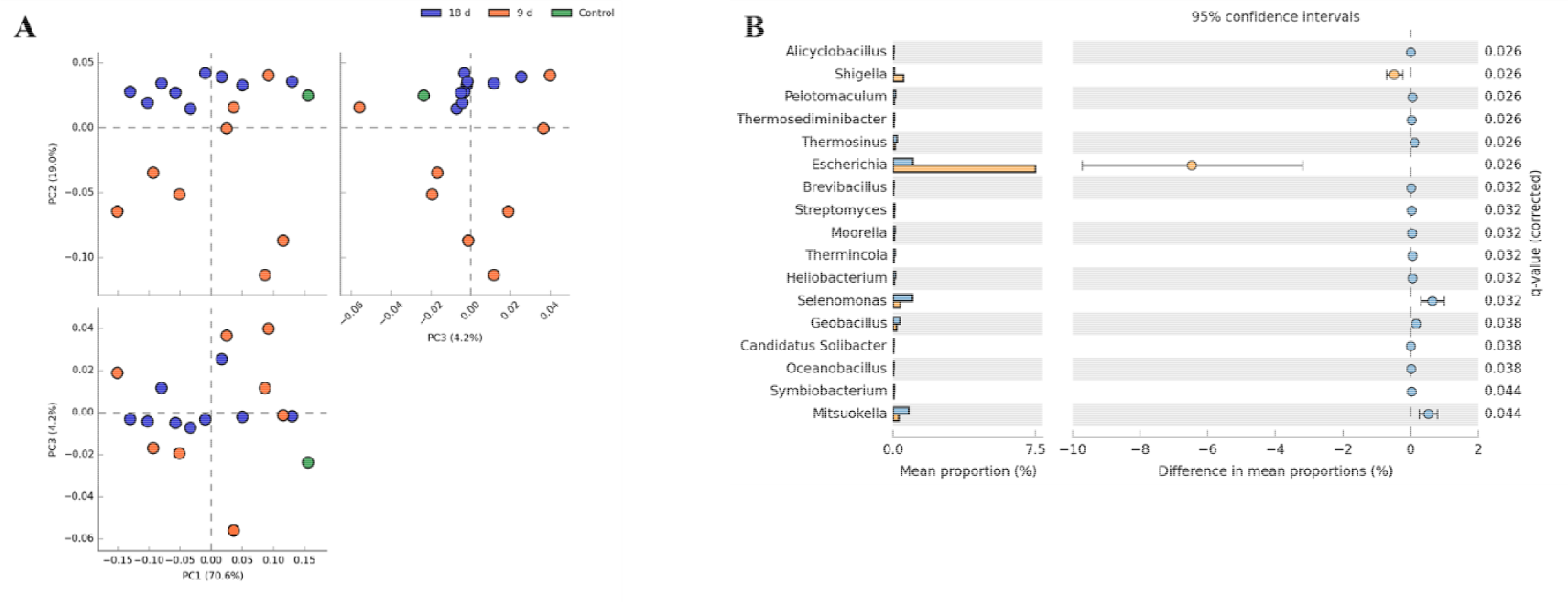
Principle coordinate analysis (PCA) of taxonomical diversity in gnotobiotic chicken. Donor material derived from 6 healthy feral chicken was orally inoculated to gnotobiotic chicken on 3 d after hatch. A) PCA analysis showed that the 18 d samples from inoculated gnotobiotic chicken was distributed closer to the inoculum when compared to 9 d samples. B) The major differences at genus level distribution were the reduced abundance of Escherichia and Shigella in the 18 d samples compared to the 9 d samples while Selenomonas, Geobacillus, and Mitsuokella increased.

Shotgun metagenomics was used to study the dynamics of the microbial community structure in the cecum of gnotobiotic chicken and the inoculum. The β-diversity represents the diversity in the compositional units between the samples. The values for Morisita-Horn index range from 0 to 1 where 1 indicates similar communities and 0 indicates dissimilar and are given in Figure 4A. All the values ranged between 0.72 and 1.0 indicating that the communities are more similar than dissimilar. However, individual variations in the colonization pattern were evident. For example, birds 4, 6 and 7 on 9 d and birds 3, 4, 6, and 7 on d 18 have dissimilar communities compared to the inoculum. However, the functional characteristics (Figure 4 B) of the communities were more similar between the samples. The range of Morisita-Horn index varied between 0.99 to 1 suggesting that the functional properties of inoculum and cecal samples at 9 d and 18 d from gnotobiotic birds were similar.

**Figure 4.**
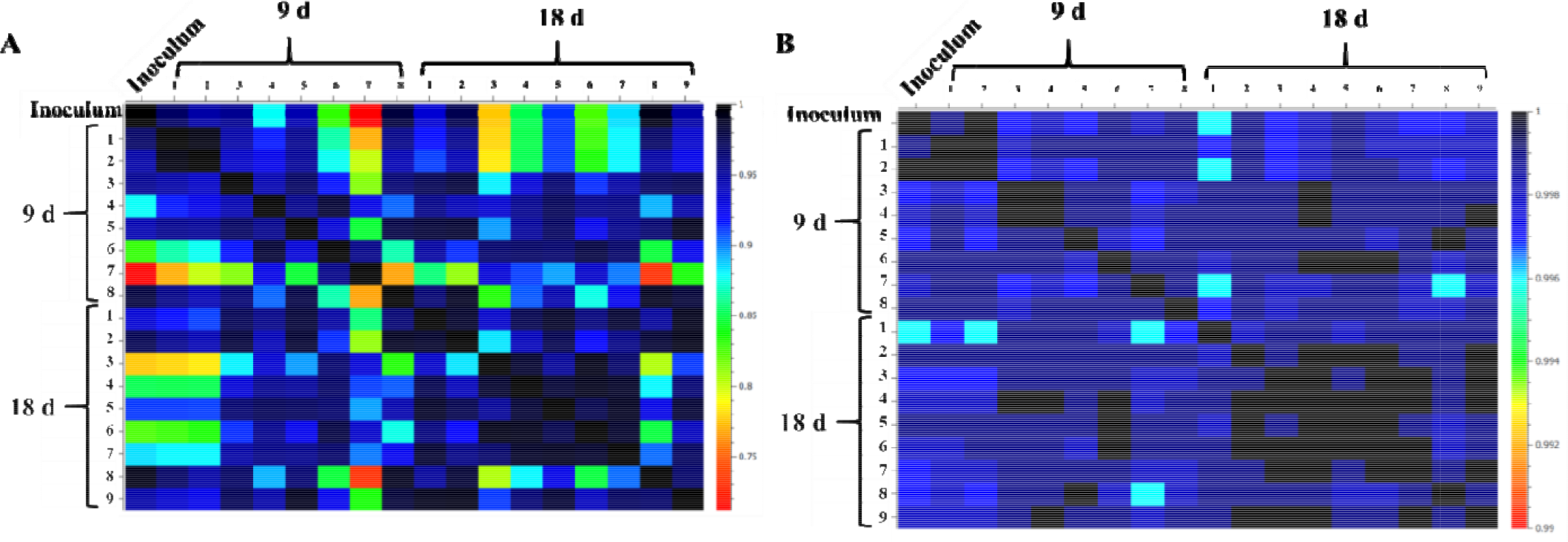
Comparison of taxonomical and functional β diversities between feral chicken-derived inoculum and gnotobiotic chicken gut samples on 9 d and 18 d. The β diversities were measured using Morisita-Horn similarity index in Explicet software. The indices range between 0 to 1 where 1 is considered similar and 0 considered dissimilar. Taxonomically, individual variations were observed between the inoculum and gnotobiotic chicken samples while the functional characteristics of the gnotobiotic chicken communities were closely similar to the inoculum.

### Functional analysis of the cecal microbiome in the gnotobiotic chicken

Analysis of functional categorization of the bacterial metagenome provides an understanding of the metabolic profile of the community. The metagenomes of feral chicken inoculum and cecal samples of gnotobiotic chicken on 9 d and 18 d were analyzed in MG-RAST pipeline using SEED subsystems database at level 2 hierarchy (Figure 5). The overall distribution pattern for ORFs that represented major cellular functions was similar for inoculum and gnotobiotic chicken microbiome on both 9 d and 18 d. In all the metagenomes, the most abundant reads (approximately 22 %) represented genes that had an unknown function. Other predominant gene function belonged to categories such as protein biosynthesis, plant-prokaryote associations, RNA processing and modification, and central carbohydrate metabolism. These findings resembled the β diversity for functional characteristics where all the communities exhibited similar profile.

**Figure 5.**
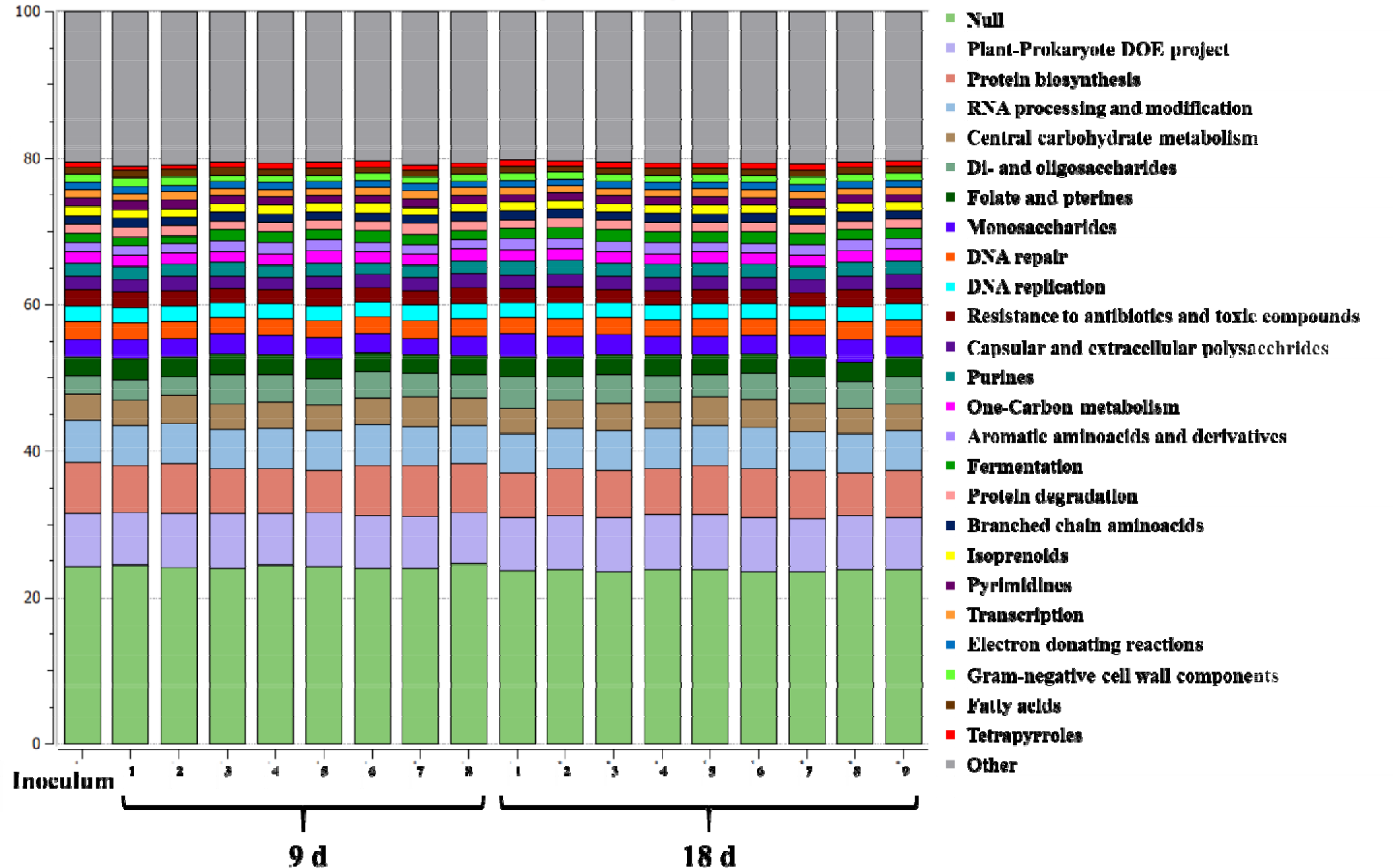
Predicted functional profile of the microbiome in feral and gnotobiotic chicken. The pooled inoculum was derived from 6 healthy feral chicken. Birds were inoculated on 3 d after hatch, were euthanized on 9 d (n = 8) and 18 d (n = 9) of age and cecal contents were collected for DNA isolation. The metagenomic functional analysis was performed in MG-RAST using Subsystems database with maximum e-value at 10^−5^ value, minimum percentage identity at 60 %, minimum alignment length of 30 amino acids and abundance 50.

## Discussion

The major objective of this experiment was to develop a gnotobiotic model to investigate the microbial succession in the cecum of gnotobiotic chickens. Various methods of rearing gnotobiotic chicken have been described previously (15, 24-27). Gnotobiotic chickens have been reared using specially designed Gustafsson germ-free apparatus (25). However, more simplified methods were developed which made use of various disinfectants to reduce the bacterial load on eggs and sterile isolators (15, 24, 26, 27). Generally, the disinfectants used were mercuric chloride, quaternary ammonium, iodoform, sodium hypochlorite solutions and commercially available chlorine dioxide solutions. In this study, Sporicidin^®^ was highly efficient in achieving disinfection without damaging the eggshells. Bacterial growth was not observed from samples collected from bird droppings and eggshells 2 d after hatching.

The conventionalization of gnotobiotic chickens using cecal microbial population derived from adult chickens has been previously conducted (15, 28). The major shortcoming of these studies was that the microbial community was identified using culture-based technique and only a few organisms could be identified. In this study, we used shotgun metagenomics to compare the microbiome of the donor material derived from apparently healthy feral chickens and the gnotobiotic chickens. By enriching for the microbial genomic DNA, shotgun metagenomics could be successfully performed using Miseq Illumina platform (29). This reduces the cost associated with using Hiseq which is the traditional platform used for shotgun metagenomics studies. Additionally, Miseq generates longer reads (250bp) compared to 150bp reads generated by Hiseq. The findings from this study indicated that gnotobiotic chicken model, when paired with next-generation sequencing techniques, could be an excellent tool to study the succession of gastrointestinal microbes in the chicken and could also be utilized in future experiments for studying the pathogenesis of enteric pathogens such as Salmonella.

The microbial population for inoculating gnotobiotic chickens in this study was collected from feral chickens that were Salmonella culture-negative. Feral chickens originated from domesticated birds that were released to the wild and have adapted to the wilderness through multiple generations. The process of feralization involves changes in social behavioral patterns, sexual selection, foraging requirements, and adaptation to predation in the wild. The expression of the genes that control these phenotypes also changes in the wild (30). Along with the host genetic changes, the microbiome could also diverge from the domesticated fowls. A study comparing the microbiome of wild and domesticated turkeys indicated that although the diversity and richness of the microbial population were similar, only 30% of the OTUs were shared between them (21). This suggested that feralization could induce adaptation of new bacterial genera to the host. Introduction of these new species to the domesticated poultry could possibly alter the microbial community in a beneficial way in the fight against enteric pathogens. In this study, we found that feral microbiome could successfully colonize in the young chicken gut without causing any detrimental effects to the health of the host.

There are few published reports on microbiome of feral chickens (Ferrario, Alessandri et al. 2017). However, the microbial population in the cecum of the pasture-housed chickens at the age of 2, 4, and 6 wk was studied recently using 16S sequencing (2). The birds were kept in conventional housing pens for first 2 wk and were released to the pasture for up to 6 wk of age. Firmicutes were the most abundant (59%) at 2 wk of age which decreased to 47% by 6 wk of age. Contrarily, the relative abundance of Bacteroidetes increased from 31% at 2 wk of age to 41% after 4 wk of pasture-housing. A substantial presence of Proteobacteria (9%) was also present in the cecal microbial population at 2 wk of age which decreased to 6% after pasture-housing. The most abundant genera in the 2 wk old chicken were Bacteroides, Ruminococaceae, and Lachnospiraceae. However, the proportion of all these genera decreased as the age progressed. Although the transient nature of microbiome with age is apparent from this study, the microbial profile may not be comparable to free-ranging feral chicken since all the birds in this experiment were fed on a formulated diet.

Conversely, the broiler chicken microbiome has been studied extensively using culture-based techniques (8, 10, 28), 16S sequencing (13, 14, 31-34), and shotgun metagenomics (6, 17). The major difference in the microbial composition of feral chicken and broiler birds was Bacteroidetes predominated in the feral chicken microbiome (66.4 %). Firmicutes represented only 18.0 % while Proteobacteria formed 9.7%. Together, these 3 phyla constituted more than 94% of the microbiome in feral chicken. On the contrary, Firmicutes were found to be the most abundant phyla in the cecum of broiler chicken. In 5 wk old broiler chickens, it was reported that 67% of the total sequences belonged to phylum Firmicutes (31). A similar finding was observed in the ceca of broiler chickens where Clostridiaceae-related sequences formed 65% of the 16S rRNA clones derived from various age groups (14). Also, other studies that analyzed the effect of age on cecal microbial composition found that Firmicutes predominated and composed more than 90% in one-week-old chicken and decreased with age to 56% by 35 d (6, 7, 35). Among the Firmicutes, Ruminococcaceae and Lachnospiraceae were found to be the most abundant families (7, 33). An age-related lowering in the abundance of Ruminococcaceae and Lachnospiraceae and an increase in the abundance of the Clostridiaceae family in the cecal microbiome was also observed recently (7). Substantial presence of Proteobacteria was also reported in the previous experiments. In 3 day-old broiler chickens, approximately 15% of total 16S rRNA sequences were identified as Proteobacteria (14), while in older chicken it formed 11 – 20 % of the total microbiome (31, 33). Proteobacteria was reported as the most abundant phylum following Firmicutes in the cecum of broiler chickens in other studies (6, 7, 35). Bacteroides was found in relatively less proportion or even absent in the cecum (6, 33). On the contrary, it was found that the proportion of Bacteroidetes increased from 2% at 15 d to 36% by 29 d of age (7). Similar to feral chicken microbiome, the presence of Actinobacteria was also detected in the cecum of broiler chicken in a recent study (35).

At the genus level, Bacteroides (52.9%) were the most abundant organisms in the feral chicken microbiome. This was followed by Clostridium (5.2%) and Prevotella (5.3%), while Ruminococcus (1.14%) and Lactobacillus (0.44%) formed lower proportions. This is in contrary to the reports from broiler chicken cecal microbiome where Ruminococcus and Lactobacilli were found to be the predominant genera. In 3 day-old chicken cecum, Ruminococcus species formed 15.6% (14) of the total sequences and in 5 week-old, it was 6% (31). Similarly, Lactobacillus species was detected at 7-8% in broiler chicken cecum in these studies. A stable proportion of 16 −23% Ruminococcus species in the total cecal microbiome which did not alter with age was observed by Ranjitkar et al. (7).

The gnotobiotic chicken microbiome from this study had different proportions of Bacteroidetes, Firmicutes, and Actinobacteria at both 9 d and 18 d compared to feral chicken inoculum. The abundance of Bacteroidetes was lower while Firmicutes and Actinobacteria were higher in conventionalized chicken compared to the feral chicken microbiome. Furthermore, in the conventionalized chickens used in this study, the proportion of phylum Bacteroidetes was relatively stable at 9 d and 18 d of age while the abundance of Firmicutes and Actinobacteria increased with age. The differences in feed and age of the birds could possibly explain these variations in the colonization profile. Dietary intervention is a primary driving force that causes alterations in the microbiome (36, 37). Feral chickens forage in the wild on a variety of feed which includes insects, berries, and worms while the gnotobiotic chicken was fed on poultry starter-diet. Another reason for the discrepancy between the feral and gnotobiotic chicken microbiome profiles could be that the inoculum was derived from pooled colon and cecal contents of feral chicken, while the analysis of gnotobiotic chicken microbiome was performed using the samples that were solely collected from the cecum.

Another finding was that although the proportion of Proteobacteria was higher at 9 d of age, it decreased by age and reached the level found in inoculum by 18 d. At the genus level, there was a decrease in the abundance of Escherichia and Shigella, which are members of Proteobacteria. This shift is analogous to the microbial succession in infant gut where initial colonization is by *Enterococcus* and *Escherichia*, followed by *Bifidobacterium* and further by obligate anaerobes belonging to Firmicutes and Bacteroidetes (38–40). Similarly, a higher proportion of Escherichia was reported in young chicken which was later substituted by obligate anaerobes (7, 35). However, the initial colonization by Proteobacteria in broiler chicken can be of public health risk especially in the context of infection by enteric pathogens such as Salmonella and Campylobacter. An early bloom in these pathogen population in broiler chicken may not be sufficiently countered by the late colonizers, thus resulting in the risk of infection even at market age (15). In this study, a decrease in abundance of Proteobacteria was correlated with an increase in Firmicutes and Actinobacteria population on 18 d. Our findings suggest that early administration of adult feral chicken microbiome could effectively prevent prolonged colonization of facultative anaerobes in chickens.

The microbial profile given in Figure 2 shows the inter-individual variation. The differences between individual birds were more pronounced at 9 d as indicated by PCA plot (Figure 3 A). Similar variation in microbial composition between the experimental birds has been reported previously (1, 2). Similar to our findings where 18 d old samples clustered together, the microbial communities from older broiler chicken clustered with less variation than communities from younger birds (35). Despite the individual variation in the microbial composition, diversity between 9 d and 18 d samples was relatively smaller. This is in discordance with previous studies where diversity was lower among the microbial population in the younger birds but increased with age (14, 35). The major reason for the increase in the diversity of microbiome with age in these experiments was that the birds were housed in pens or cages and could acquire newer organisms from the habitat. In this experiment, the inoculum served as the sole source of microbes and successful colonization of this microbiome happened by 9 d of age.

Alternatively, the functional properties of the microbial communities were more stable at 9 d and 18 d and were similar to the feral chicken inoculum even while the microbial composition was different. There were no significant differences between the inoculum, 9 d, and 18 d samples for the functional properties even at level 2 hierarchy using SEED Subsystems database. Similar results for functional properties of chicken cecal microbiome was observed previously (6, 17, 35). The variability between individuals for the taxonomic profile occurring during the microbial succession did not reflect in the functional profiles in these studies. It has been found that the microbes occupying equivalent niches share similar functional properties even in diverse hosts (41). The microbial assemblage in a previously uninhabited habitat could be driven by equivalence in functional aspects rather than the stochastic nature of microbial colonization. In this study, the host niches being similar, the evenness in functional properties of the communities despite taxonomical variability could only be explained if functionally similar organisms are occupying the equivalent niches.

Chickens act as a reservoir for enteric human pathogens especially, Salmonella. Recently, various serotypes such as Enteritidis, I,[5],12:i:-; Typhimurium; Heidelberg; Hadar; Mbandaka; Montevideo; Agona; and Infantis has been associated with Salmonella outbreaks (42, 43). These rampant multistate Salmonella outbreaks due to transmission from live poultry reveal the necessity of pursuing studies aimed at control of Salmonella in poultry. The presence of enteric pathogens in poultry was controlled by using antibiotic feed additives (44). Due to the recent FDA regulations to limit the use of antimicrobials in the food supply due to public health concerns, use of such antibiotic feed additives is currently highly controlled. It is pertinent to develop alternatives such as prebiotics and probiotics that could manipulate the microbial community in chicken and thus competitively exclude enteric pathogens. The pioneering work by Nurumi and Rantala in 1973 (10) demonstrated the competitive exclusion of Salmonella by adult chicken microbiome while microbiome from young birds was incapable to prevent the growth of Salmonella (15). The recent outbreaks suggest that this subject should get renewed attention as there are evidence for more Salmonella serotypes adapting to chickens and causing a potential threat to public health (42). Gnotobiotic chickens could serve as excellent models for studying the microbial colonization resistance towards these pathogens and also could be used for development of probiotics as alternatives for controlling the pathogens.

## Materials and Methods

### Experiment and sampling

Feral chickens were obtained locally near Brookings, South Dakota, USA. The feral flock was once a captive flock of mixed breed and has been feral for no less than 8 years. Birds forage on a small grain farm and in surrounding grasslands. Feral chickens were sampled during a routine slaughter for personal meat consumption by the land owners. Gut samples from six birds were collected from the visera following slaughter. The intestine was ligated at distal ileum and distal colon, maintained in ice, and transported immediately to the laboratory and stored at −80C. Protocols used in this study for sample collection were reviewed and approved by the Institutional Animal Care and Use Committee (IACUC) at the South Dakota State University, Brookings, South Dakota. For processing, samples were transferred to a Coy Anaerobic chamber and the contents were expelled into 50 ml sterile conical tubes. Samples were diluted 1:10 (w/v) using anaerobic Brain Heart Infusion broth supplemented with volatile fatty acids and vitamins (BHI-M), mixed by repeated pipetting, and aliquoted into cryovials. Anaerobic DMSO was added at 18% (final conc) and stored at −80°C until inoculation into young chickens. Simultaneously, an aliquot of the, neat samples from the intestines were streaked on Xylose Lysine Tergitol-4 (XLT-4) agar plates and incubated aerobically at 37°C for 24 h. For preparing inoculant into germ-free chickens, samples were thawed and stock from 6 feral chickens was pooled at equal volume and further diluted 1:10 using anaerobic PBS.

Gnotobiotic chickens were reared using a modified protocol that was described previously (15). Eggs of White Leghorn chickens were acquired from a commercial hatchery, treated with Sporicidin^®^ disinfectant solution (Contec ^®^, Inc.), with sterile water and incubated in an incubator, pre-treated with Sporicidin^®^, at 37 °C at 55% humidity. Humidity was maintained using a 1% (wt/vol) aqueous solution of potassium permanganate. After 19 d of incubation, eggs were removed from the incubator and candled for viability. Viable eggs were transferred to an UV biosafety cabinet and dipped in Sporicidin^®^ solution for 15s then wiped with a sterile cloth saturated with sterile water. Eggs were then transferred to autoclaved egg trays, placed in sterile autoclave bags and transferred immediately to the port of the isolator unit. Eggs were sprayed with 5% peracetic acid, and after 20 min exposure, were transferred inside the isolators. Eggs were maintained at 37°C at 65% humidity until hatching on 21 d.

Following hatching, birds were provided *ad libitum* sterilized water and gamma-irradiated starter diet (LabDiet^®^ 5065, Irradiated) designed to meet nutrient requirements of young chickens (Table 1) and monitored daily. On 3 d post-hatch, birds (n=17) were inoculated orally with 300 µL of pooled cecal contents. Eight birds were euthanized using cervical dislocation on 9 d and 9 birds were euthanized on 18 d post-hatch. The cecal contents were collected for DNA isolation and stored at −20°C until use.

**Table 1.** Nutritional composition and energy content of the LabDiet^®^ 5065, Irradiated diet.

| **Composition** | **in %** |
| --- | --- |
| Protein | 22.1 |
| Fat (ether extract) | 4.2 |
| Fat (acid hydrolysis) | 5.2 |
| Fiber (max) | 2.8 |
| Nitrogen-Free Extract | 55.6 |
| Minerals | 5.3 |
| **Energy source** |  |
| Protein | 25.4 |
| Fat (ether extract) | 10.7 |
| Carbohydrates | 63.9 |
| Total Energy (kcal/g) | 3.48 |

To assess the sterility of the isolator, swabs were collected on 2 d post-hatch from the egg shells, droppings, and isolator floor and transferred to anaerobic transport media (45) and removed from the isolator. The swabs were then streaked on BHI-M agar plates and incubated aerobically at 37°C. The plates were examined for the presence of bacterial colonies after 24 h and 48 h of incubation.

### Genomic DNA isolation from the cecal contents

Genomic DNA was isolated using Powersoil DNA isolation kit (Mo Bio Laboratories Inc, CA). Briefly, approximately 100 mg of cecal contents were transferred to bead tubes and samples were homogenized for 2 min using TissueLyser (Qiagen, Germantown, MD). DNA isolation was performed according to manufacturer’s protocol and DNA was eluted in 50 µL nuclease-free water. The quality of genomic DNA samples was assessed using NanoDrop™ One (Thermo Fisher Scientific, Wilmington, DE) and quantified using Qubit Fluorometer 3.0 (Invitrogen, Carlsbad, CA). Samples were stored at −20°C until use.

### Microbial DNA enrichment and shotgun metagenomics sequencing

Selective enrichment of bacterial genomic DNA was performed using NEBNext^®^ Microbiome DNA Enrichment Kit (New England Biolabs, Inc. MA) following methods previously published by our group(29). Briefly, 0.5 µg of genomic DNA was treated with 80 µl of MBD2-Fc bound magnetic beads in the presence of binding buffer and incubated at room temperature for 15 min with rotation. After incubation, beads were separated by keeping the tubes on a magnetic rack for 5 min. The supernatant containing microbial DNA was transferred to a fresh tube. The DNA was further purified using Agencourt AMPure XP beads (Beckman Coulter) and stored at −20°C.

For shotgun metagenome sequencing, the enriched genomic DNA from pooled feral samples, 8 samples from 9 d and 9 samples from 18 d post-hatch gnotobiotic chickens was used. The concentrations of genomic DNA samples were adjusted to 0.3 ng/µl. Samples were then processed using Nextera XT DNA Sample Prep Kit (Illumina Inc. San Diego, CA) according to manufacturer’s protocol. Purified products with unique barcodes were normalized using bead normalization protocol of the manufacturer and equal volumes of normalized libraries were pooled together and diluted in hybridization buffer. The diluted libraries were heat denatured prior to loading to the sequencer. Illumina paired-end sequencing was performed on the Miseq platform using a 2 − 250 paired-end sequencing chemistry.

### Sequence data processing

The raw data files were de-multiplexed and converted to fastq files using Casava v.1.8.2. (Illumina, Inc, San Diego, CA, USA). Fastq files were concatenated and analyzed using MGRAST pipeline (46). The quality control steps in MG-RAST included dereplication, ambiguous base filtering, and length filtering. The taxonomical abundance was analyzed using MG-RAST with Best Hit Classification approach using Refseq database and parameters were limited to minimum e-value of 10^−5^, minimum percentage identity of 60%, a minimum abundance of 50, and a minimum alignment length of 30 amino acids. The functional abundance was analyzed using Hierarchical Classification in MG-RAST using SEED Subsystems database and parameters were limited to minimum e-value of 10^−5^, minimum percentage identity of 60%, a minimum abundance of 50, and a minimum alignment length of 30 amino acids. The OTU abundance tables were downloaded from MG-RAST and were used for downstream statistical analysis.

### Statistical analysis

The beta diversity between the feral chicken inoculum, 9 d, and 18 d samples was estimated using Morisita-Horn index in Explicet software (47). The PCA analysis for taxonomical and functional diversity was performed using STAMP (48). Also, the differences in genus-level distribution between 9 d and 18 d samples were calculated in STAMP software using White’s non-parametric t-test with Storey false discovery rate correction and filtered for a minimum of 200 reads. The differences in phylum-level distribution between 9 d and 18 d samples were calculated in GraphPad prism 7.03 (GraphPad Software, San Diego, CA) using non-parametric Mann-Whitney U test. A significant difference was recorded at P value < 0.05. Genera-level distribution tables were analyzed using Explicet software.

## Author Contributions

JS conceived and designed the experiments. MT, SW, SG, and AS performed the experiments. MT analyzed the data. JS, KD, EN, SC, TW and TJ contributed reagents and materials. MT, JS and KD wrote the manuscript. All authors reviewed and approved the manuscript.

